# A computational model of the mammalian auditory periphery with a multichannel, energy-driven, medial olivocochlear reflex

**DOI:** 10.1101/2025.11.26.690762

**Authors:** Daniel R. Guest, Afagh Farhadi, Laurel H. Carney

## Abstract

The afferent auditory system, and how its specialized mechanisms and circuits support ecologically relevant auditory computations such as speech recognition, has received considerable attention in decades past. This work has culminated in accurate computational models of early afferent coding alongside a good understanding of how low-level mechanisms (e.g., peripheral tuning) impact auditory perception. In contrast, the auditory efferent system and its role in auditory perception is much less well understood. To address this gap in knowledge, we describe modifications to a model of the auditory periphery to include a medial olivocochlear efferent pathway that dynamically adjusts cochlear gain in response to sound via the classical medial olivocochlear reflex loop. We show that this model can simulate the effects of contralateral elicitors on auditory-nerve responses, including the effect of elicitors that are tonotopically distant from probes. Inclusion of across-frequency efferent effects necessitated a novel multichannel design.

## I. INTRODUCTION

The auditory efferent system comprises descending connections from higher-to lower-level auditory brain areas and the auditory periphery (Guinan, 2018; Ryugo and Fay, 2010). These include connections from cortical to subcortical neurons, from subcortical to other subcortical neurons, and from subcortical neurons to various targets in the cochlea, such as outer hair cells (OHCs) and auditory-nerve fibers (ANs). This final set of circuits—the olivocochlear (OC) system—is of principal interest here. The OC system consists of neurons located in or nearby the superior olivary complex that send projections to targets in the cochlea. In mammals, the OC system is differentiated into two sub-pathways, named for the positions of their constituent cell bodies relative to the superior olivary complex: the medial olivocochlear (MOC) system and the lateral olivocochlear (LOC) system (Guinan, 2018; Warr, 1992). We defer further consideration of the LOC system to Sec. IV and focus our attention on the MOC system.

MOC neurons receive excitatory ascending input from the cochlear nucleus (Brown et al., 2003, 2013; Darrow et al., 2012) and send myelinated projections to the cochlea to form synapses on OHCs (Warr and Guinan, 1979), Type I AN dendrites (Hua et al., 2021), and Type II AN dendrites (Bachman et al., 2025). The direct influence of MOC neurons on OHCs, which is the focus here, is much better understand than their direct influence on AN fibers, for which no physiological data is currently available. When MOC neurons are stimulated, they release acetylcholine, hyperpolarizing OHCs via α9α10 cholinergic receptors and reducing cochlear gain (Wersinger and Fuchs, 2011). The consequences of this gain reduction have been observed invasively in basilar-membrane vibration (Cooper and Guinan, 2006; Dolan et al., 1997), inner-hair-cell potentials (Brown et al., 1983; Brown and Nuttall, 1984), and afferent-fiber activity (Guinan and Gifford, 1988a, 1988b, 1988c; Guinan and Stankovic, 1996; Warren and Liberman, 1989a, 1989b), observed non-invasively via otoacoustic emissions (Lilaonitkul and Guinan, 2009; Walsh et al., 2010; Wojtczak et al., 2015, 2019) and psychophysics (Almishaal et al., 2017; Jennings, 2021; Jennings et al., 2011, 2018; Roverud and Strickland, 2015; Strickland, 2004, 2008; Yasin et al., 2014), and simulated via computational models (Farhadi et al., 2023; Grange et al., 2022; Messing et al., 2009; Smalt et al., 2014; Yasin et al., 2018). Because MOC neurons respond to sound, increase their firing rate with increasing sound level, and reduce cochlear gain (Brown, 1989; Brown et al., 2003), the MOC system has often been described as a sound-driven “reflex”, and there is great curiosity about the possible role(s) of this reflex in auditory perception (Jennings, 2021).

In this paper, we present a new auditory computational model that includes a brainstem-level MOCR circuit. To motivate the design of our MOC model, we first highlight several key anatomical characteristics and physiological characteristics of the MOC system. The number of MOC neurons varies significantly among species (with as few as ∼350 in hamster and as many as ∼2400 in guinea pig) but is much smaller than the number of innervated OHCs; each MOC neuron typically innervates 20–80 OHCs, and each OHC receives input from 1–2 MOC neurons on average (Brown, 2011, 2014; Liberman and Brown, 1986; Ryugo and Fay, 2010; Steenken et al., 2024; Warr, 1992). Individual MOC neurons tend to synapse onto OHCs tuned to similar frequencies, but MOC neurons branch in the cochlea, innervating OHCs spanning 5-10% of cochlear distance on average, but sometimes as little as 1% or as much as 25% of cochlear distance (Brown, 1989, 2014; Liberman and Brown, 1986). This synaptic patterning is depicted schematically in Fig. 1A. MOC neurons respond to sound, exhibit “chopping” type responses, have sharp frequency tuning (comparable to or slightly broader than afferent tuning, depending on the species and study), have dynamic ranges exceeding 40 dB for pure-tone stimulation, and adapt weakly compared to AN fibers (Brown, 1989, 2001; Brown et al., 2003; Liberman and Brown, 1986). Limited data suggests that MOC neurons are also tuned to amplitude-modulation frequency (Gummer et al., 1988), but generally little is known about how they respond to other higher order features of sound.

**Figure 1.**
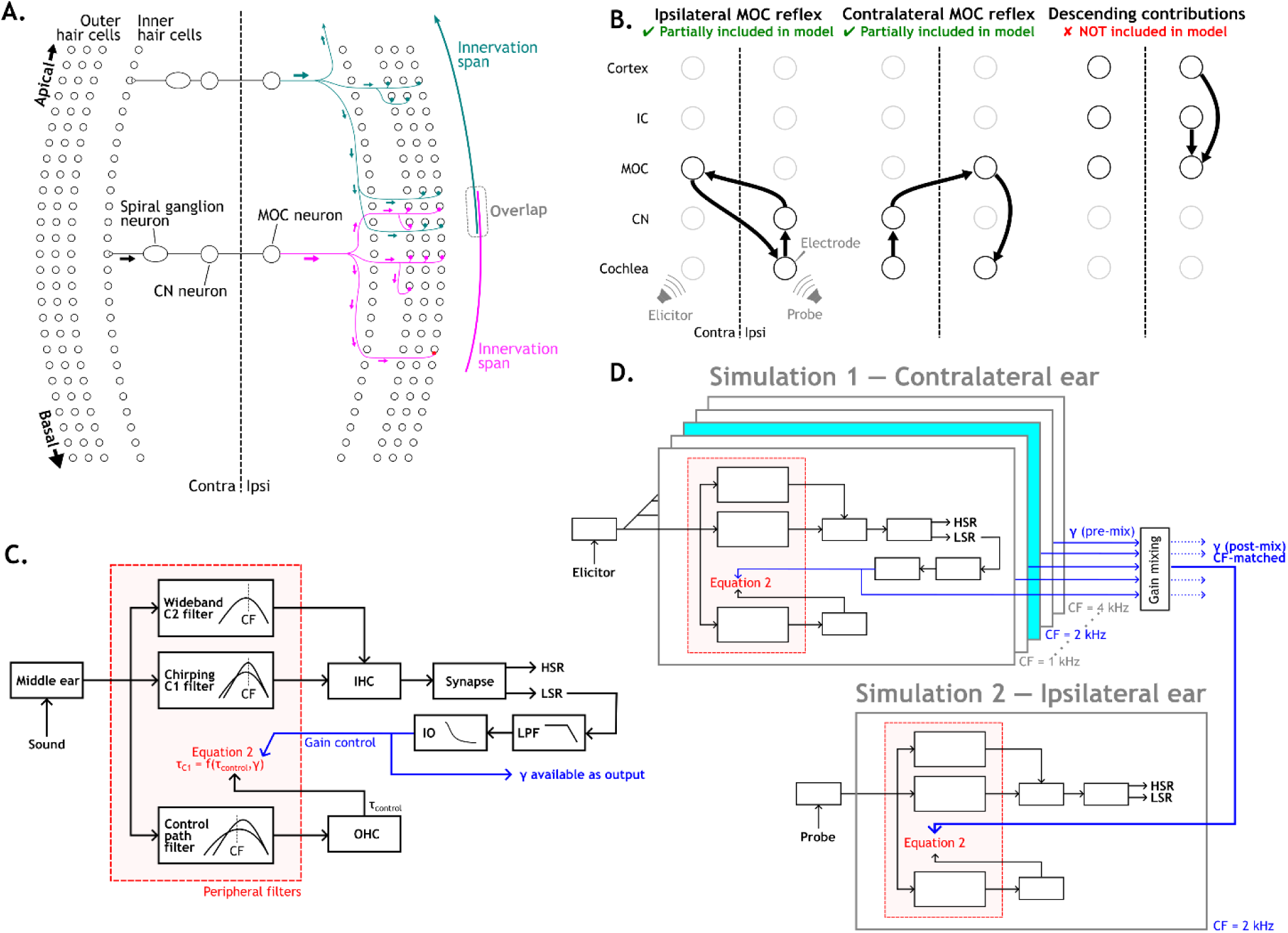
(A) Schematic depicting the conceptual model underlying the present simulation of the contralateral MOCR. On the contralateral side (left), an elicitor drives activity in AN fibers, which subsequently excite tonotopically similar cochlear nucleus neurons located on the same side as the elicitor and, after a midline cross, tonotopically similar MOC neurons on the opposite side as the elicitor. These MOC neurons each then project to innervate multiple ipsilateral OHCs. Crucially, because of the sometimes wide tonotopic span of innervation patterns by individual MOC neurons (Brown, 2014), cochlear gain at an ipsilateral (right) recording site could be influenced by the activity of multiple MOC neurons tuned to different frequencies. (B) A series of schematics indicating contributions to MOC activity that could influence cochlear gain in the ipsilateral cochlea. From left to right: the ipsilateral MOCR, the contralateral MOCR, and descending inputs to MOC. (C) Schematic depicting a single channel in the model with MOCR (c.f. Fig 1, Zilany and Bruce, 2006). Blue arrows indicate the path of the control signal for the MOCR (γ), which is used internally but also provided as an output for subsequent analysis or simulations. (D) Schematic depicting how CAS was approximate via two separate single-ear simulations (Simulation 1 and Simulation 2). First, a population response to the contralateral elicitor stimulus is simulated (Simulation 1, top). Then, a single-channel response to the ipsilateral probe stimulus is simulated at the CF of interest (Simulation 2, bottom). However, instead of allowing cochlear gain to be dynamically determined based on the MOCR (as in the single channel depicted in C), the MOCR control parameter (γ) is extracted from the outputs of Simulation 1 and used to determine cochlear gain at all time points during Simulation 2.

An important aspect of the MOC system is its binaural structure. Most MOC neurons can be characterized as falling into one of two stereotyped circuits; the ipsilateral reflex circuit or the contralateral reflex circuit (Guinan, 2006). The brainstem-level ipsilateral reflex comprises three connections, from (1) ipsilateral AN fibers to ipsilateral CN neurons (2) to contralateral MOC neurons (3) to ipsilateral OHCs (i.e., “double crossed”; Fig. 1B, left). The contralateral reflex also comprises three connections, from (1) AN fibers to contralateral CN neurons (2) to ipsilateral MOC neurons (3) to ipsilateral OHCs (i.e., “single crossed”; Fig. 1B, middle). The anatomy of these circuits is consistent with observations that most MOC neurons respond preferentially to sound presented to one side of the head, and that few MOC neurons innervate both cochleae (Kishan et al., 2011). In lab mammals, the ipsilateral reflex appears to be stronger than the contralateral reflex (Guinan and Gifford, 1988a). In humans, similar results are reported for narrowband stimuli, but for broadband stimuli ipsilateral and contralateral reflexes appear to be more similar in strength (Lilaonitkul and Guinan, 2009). It is important to note that these brainstem-level reflex circuits constitute only a subset of the full circuits that govern MOC gain control, as MOC neurons also receive descending input from inferior colliculus (Romero and Trussell, 2021, 2022) and auditory cortex (Doucet et al., 2002) (Fig. 1B, right). The issue of other sources of input to MOC is further discussed in Sec. IV.

Many strategies have been tested to shed light on MOC function, varying primarily in the provenance of the data—mechanical, otoacoustic, neural, or psychophysical—and in the strategy used to elicit the MOCR, such as electrical stimulation of MOC fibers (Brown et al., 1983; Brown and Nuttall, 1984; Elgueda et al., 2011; Guinan and Gifford, 1988a, 1988b, 1988c; Guinan and Stankovic, 1996), contralateral acoustic stimulation (CAS) (Faubion et al., 2024; Kawase et al., 1993; Warren and Liberman, 1989a, 1989b), or elicitor-probe designs (Jennings et al., 2011; Roverud and Strickland, 2015; Strickland, 2004, 2008). In developing the present model, we narrowed our focus to data from single-unit AN recordings that employed CAS to elicit the MOCR, a strategy guided by several lines of reasoning. First, CAS is more naturalistic and straightforward to model than electrical stimulation, and it is easier to generalize from CAS responses to other sound-driven responses than from electrically driven responses to sound-driven responses. Second, the baseline auditory model that we modified (Zilany et al., 2014) focuses on simulating AN responses, and thus AN responses are the most natural point of contact between model and data for our purposes. Finally, the key features of AN-CAS data are fairly straightforward and consistent with data from other modalities. Within 100–200 ms of MOC activation by a contralateral elicitor, AN fibers show elevated thresholds, rightward shifts of rate-level functions (RLF) for tones presented near the fiber’s characteristic frequency (CF), and relatively little change in response to tones at other frequencies (Warren and Liberman, 1989a, 1989b). These effects are largest in the mid-CF region (near 2–4 kHz in cat), require contralateral elicitors to be presented above a threshold level, and are consistent with MOC-driven changes in cochlear gain being the primary mechanistic factor at play.

Here, we modified a previously published auditory model (Zilany et al., 2014) to include the MOCR, calibrating the model architecture and parameters manually to qualitatively match trends in existing AN-CAS data (Warren and Liberman, 1989a, 1989b). Our model differs from previous MOCR models in a few key ways. First, unlike our own lab’s previous MOCR model that also included descending inputs from the inferior colliculus to MOC neurons (Farhadi et al., 2023), the present model focuses only on the brainstem-level circuits thought to be driven primarily by stimulus energy. Second, unlike prior models of the MOCR that regulated the gain of each channel based only on afferent responses from the same channel (single-channel MOCR; e.g., Smalt et al., 2014), the present model employs a multichannel design that allows afferent responses to impact cochlear gain in tonotopically distant channels. The model MOCR was driven by lowpass-filtered low-spontaneous-rate (LSR) responses, which served as a proxy for wide-dynamic-range level-sensitive neurons in the auditory brainstem that are thought to drive the efferent reflex (Farhadi et al., 2023). CAS was not simulated directly (i.e., with a true binaural MOC model) but using an approximate strategy that simplified implementation. The resulting model innovates on existing models in the literature in several ways and maintains reasonable computational costs despite the significant increase in model complexity. We anticipate that this model will help to identify non-invasive behavioral and physiological correlates of MOC-driven cochlear gain control.

## II. METHODS

### A. Model algorithm

The present computational model was adapted from an existing computational model described in Zilany et al. (2014) in a manner inspired by Farhadi et al. (2023). Earlier model versions were single-channel models broken into two separate time loops: one loop for simulating IHC responses given an input time-pressure waveform and a second for simulating AN responses given an input IHC voltage waveform Responses from multiple independent channels were generated by repeated calls to the model functions in a loop. The present model, in contrast, is a multi-channel model with a single time loop. That is, each step of the time loop receives a single sample of an input time-pressure waveform and returns a single output sample from every model stage in every model channel. This change allowed responses from any stage or channel to, in theory, serve as control signals to affect parameters in any other model stage or channel. The model time loop was calculated in the following way:

1. Before a simulation began, the input sound-pressure waveform was prepended with 20 ms of zeros. This time was needed in practice to allow several stages in the model time to stabilize before simulation of sound-driven responses. A similar system was used in previous versions of this model but was CF-specific, with longer stabilization times for low-CF channels and shorter stabilization times for high-CF channels. In the present model, for ease of implementation, the same 20-ms value is used for all channels regardless of CF. At the end of a simulation, this additional time was removed before returning the outputs.
2. As in prior versions of the model, the input sound-pressure waveform was passed through a species-specific middle-ear filter (either human or cat) and delayed in time. The time-delay values used were the same as in Zilany et al. (2014), which were intended to match simulated reverse-correlation functions to empirical reverse-correlation functions (Zilany and Bruce, 2006).
3. Delayed middle-ear outputs were passed through the peripheral filtering stage used in Zilany et al. (2014). In earlier versions of the peripheral filter model, the level-dependent tuning and gain of the “C1” signal-path filter was determined by manipulating the filter time constant based on responses to sound from the parallel “C2” control-path filter and a scalar parameter ***C***_OHC_. Specifically, the C1 filter time constant was set to

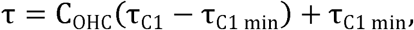

where τ_C1_ is the current C1 time constant (as determined by responses from the C2 filter) and τ_C1 min_ is the shortest possible time constant^1^ (Zilany and Bruce, 2006). *C*_OHC_, which varies between 0 and 1, can be understood as interpolating between a “normal-hearing” state for ***C***_OHC_ = 1, corresponding to the longer time constant τ_C1_, a narrower bandwidth, and more gain, and an impaired state for ***C***_OHC_ = 0, corresponding to a shorter time constant of τ_C1 min_, a broader bandwidth, and less gain. In the present model, we built atop this system replacing the equation for τ above with

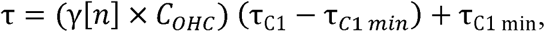

where γ[n] is a time-varying value in (0, 1) that is determined by model responses in the MOC stage (see below). This calculation of τ allows responses in the MOC stage to influence cochlear gain at subsequent time steps in a dynamic fashion.
4. As in prior versions of the model, outputs from the peripheral filtering stage were then passed through a simulation of inner-hair-cell (IHC) transduction, adaptation in the IHC-AN synapse complex, and absolute refractoriness (Zilany et al., 2009). In the original model code, some of these steps used different parameter values to simulate responses from either high-spontaneous-rate (HSR) or LSR neurons. We made three notable changes from the previous model in this stage. First, IHC potentials were *not* downsampled to 10 kHz before subsequent processing, to avoid the complexities associated with operating different parts of the model at different sampling rates. Second, power-law adaptation was implemented with the approximation described in Guest and Carney (2024) instead of directly or using previous approximation strategies. Third, responses from both HSR and LSR fibers were simulated in a single function call, rather than the user to choosing one or the other via an input argument. This modification allowed HSR model responses to depend on the response history of LSR fibers (see below).
5. To simulate MOC gain control, LSR rates were first lowpass filtered with a single-pole IIR lowpass filter (***f**_c_* = 0.64 Hz) to account for the sluggish temporal dynamics of the MOCR (Backus and Guinan, 2006; Warren and Liberman, 1989a).
6. Next, filtered LSR rates were subjected to a static nonlinearity to map from rate in [0, ∞) to a value in [0, 1] that we refer to as a “gain control factor” or γ, according to:

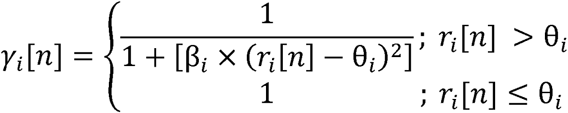

where γ_*i*_[*n*] is the gain-control factor for the *i*^th^ frequency channel at time step ***n, r_i_***[*n*], is the lowpass-filtered LSR rate in the *i*^th^ frequency channel at time step *n*, β_*i*_ is a parameter governing the slope between rate and gain-control factor in the *i*^th^ frequency channel (IO slope), and θ_*i*_ is a parameter (in sp/s) that determines the rate that must be exceeded before gain reduction can take place in the *i*^th^ frequency channel (IO threshold). This nonlinearity matched the nonlinearity used in the analogous stage of the MOC model of Farhadi et al. (2023), albeit with different parameter values. To account for differences suggested by AN data in the MOCR threshold and magnitude across CF (see Sec. III), we heuristically adjusted the values of these parameters across CF according to:

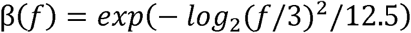

and

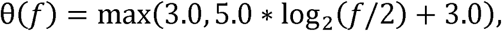

where ***f*** is the CF in kHz.
7. Next, gain control factors were mixed across channels using a Gaussian weighting function with a standard deviation of one octave (and normalized to integrate to 1, like a Gaussian probability density function) to account for the broad innervation of OHCs by MOC neurons in the cochlea (Brown, 2014). That is, each channel’s “post-mix” gain factor was a weighted sum of all channels’ “pre-mix” gain factors from the same timestep, where the weighting function was a normalized Gaussian defined on a log-CF scale and centered on the channel’s CF. Note that, because gain-control factors are limited to the range [0, 1], this mixing was performed using a weighted geometric average rather than a weighted arithmetic average. (Equivalently, we could have mixed log-transformed gain factors and then exponentiated the result.)
8. Finally, MOC responses were delayed in time by 25 ms to account for the latency of MOC effects in the cochlea (Backus and Guinan, 2006).

### B. Acquisition and analysis of physiological data

Physiological data were extracted from published figures in Warren and Liberman (1989a,b) using WebPlotDigitizer (Rohatgi, n.d.). In several cases, many overlapping points were present in individual figures and it was difficult to tell how many data points to mark. Nevertheless, because the data were used only to estimate trendlines in the data, the quality of the extracted data were sufficient for our purposes. A correction was made for axis rotation because the cited articles are scans of original printed copies, rather than original digital copies, and are not perfectly oriented with respect to the page. These data were collected into tidy-format CSV files and made available alongside our code online (see Data Availability).

We estimated trendlines from the physiological data to summarize general patterns in the context of high variance in data pooled across animals and units. To do so, a second-order polynomial was fit to the data using Optim.jl (Mogensen and Riseth, 2018) using Newton’s method via forward-mode automatic differentiation. This optimization was either performed with a standard mean-square-error (MSE) loss function, to estimate a trendline through the local mean of the data, or with the custom composite loss function

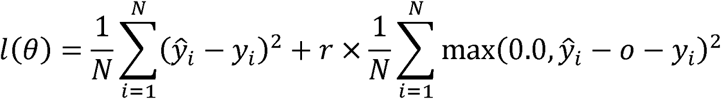

to estimate a trendline along the lower envelope of the data (a mirrored loss function was used to (left) and an aggressive penalty (with hyperparameter *r* = 200) against predictions that fell above estimate the upper envelope). This loss function balanced between finding a fit that minimized MSE envelope of the data, although on occasion the additional hyperparameter *o* was adjusted to the data (right). In practice, this loss function yielded qualitatively good estimates of the lower introduce a further offset between data and envelope. Because the envelope trendline was never used to perform analyses (only to provide a convenient visual reference on figures), adjustment of the hyperparameter *o* was performed manually to improve visual clarity in the final figures.

### C. Simulation and stimulus methods

Stimuli were synthesized using custom software in Julia 1.11 at a sampling rate of 100 kHz. Stimuli consisted of either pure tones or Gaussian noise in various binaural configurations and temporal configurations which approximated those used in the AN experiments; see below for details (Warren and Liberman, 1989a, b). Except where otherwise specified, stimuli were ramped on and off with 5-ms raised-cosine windows, with stated durations including the ramped portions. Probe (ipsilateral^2^) stimuli were always pure tones. Elicitor (contralateral) stimuli were most often pure tones, but in some cases, indicated in the text, were broadband or filtered Gaussian noise. Further stimulus details are presented in the subsections below for each type of simulation conducted.

All simulations were carried out with the updated model algorithm and parameter values described above using the Julia version of the wrapper code (see Data Availability). All other parameter values matched their default values for the cat version of the AN model from Zilany et al. (2014). The model sampling rate was set to 100 kHz. To focus on underlying trends in model responses free of stochastic components, the instantaneous rate output was used instead of the spiking output, and fractional Gaussian noise in the adaptation stage of the model was disabled (i.e., input noise values were set to zero at all time steps) and the spontaneous-rate parameter was set to a fixed value based on Zilany et al. (2014). Although we disabled fractional Gaussian noise for the present simulations, users who wish to simulate responses with fractional Gaussian noise enabled for other purposes can still do so in the provided model code. When computing average rates, only rates from times at least 50-ms after probe onset were included to minimize the contribution of onset responses to average rate, and offset responses were excluded.

As mentioned briefly above, CAS was not simulated using a “true” binaural MOC model. Instead, the effect of CAS at a given ipsilateral CF of interest was simulated using an approximation strategy. First, a population response model was simulated in response to the contralateral component of the binaural stimulus (Figure 1D). Unless otherwise noted, this simulation was carried out for 21 CFs spaced logarithmically around the ipsilateral CF of interest. Then, the spatially mixed time-varying gain-control factor from the contralateral channel tuned to the same CF as the ipsilateral channel of interest was extracted and used as the gain-control factor (γ) during subsequent simulation of the response at CF to the ipsilateral stimulus. During the simulation of the ipsilateral response, dynamic MOC gain control was disabled, and the gain-factor at each time step was instead determined by gain-control factor extracted from the contralateral side. This strategy had two key advantages. First, it eliminated the need to modify the source code of the model to simulate both a left and right ear. Second, it allowed us to pay the cost for simulating a population model only once for the contralateral ear, where responses in many channels were hypothesized to contribute to the overall result due to the spatial-mixing step in the gain-control calculations, but not for the ipsilateral ear, where only the response at CF was of interest. From a scientific standpoint, this strategy was justified because the primary configuration tested in our simulations involved relatively low-level ipsilateral probes being affected by relatively high-level contralateral elicitors (see further discussion of this issue in Sec. IV).

#### 1. Rate-level functions

RLFs involved simulating responses to 100-ms ipsilateral pure-tone probes with 5-ms raised-cosine ramps over a range of sound levels and computing the average rate within the last 50-ms of the probe. Typically, levels from 10 to 70 dB SPL were tested in steps of 2.5 dB, except in some cases where lower levels were needed to ensure that threshold was visible in the RLF. Following simulation, the rate-level curve was then interpolated by optimizing a logistic function to fit the data using the same optimization algorithm described in Sec. II.B. and then sampling the fitted function at 1000 points between the lowest and highest simulated levels. Subsequent analysis of RLFs (e.g., determination of midpoint, see Sec. III), or comparisons between RLFs. was always performed on the interpolated RLFs. In some cases, RLFs were simulated with a contralateral elicitor. In these cases, the parameters of the elicitor are reported in the text, but generally elicitors were 1000-ms stimuli with 5-ms raised-cosine ramps gated on 900 ms before the probe and gated off at the same time as the probe (Fig. 2). This temporal configuration allowed the MOCR sufficient time to build up to steady state before measuring its effects via the probe response. Although not identical, this temporal configuration should emulate the configuration of Warren and Liberman (1989a), combining long-duration precursors with short probes. Note that even when a contralateral elicitor was not present or the when the model used was an ipsilateral-only single-channel control model, the temporal configuration of the stimulus was maintained to match other simulations; i.e., the probe was always presented 900 ms after simulation start, as it would have been if a contralateral elicitor were present. This match across conditions ensured that differences between measured RLFs were always due to the properties of the elicitor, rather than differences in the amount of time simulated (leading to, for example, differences in the degree of adaptation in the nerve response).

**Figure 2.**
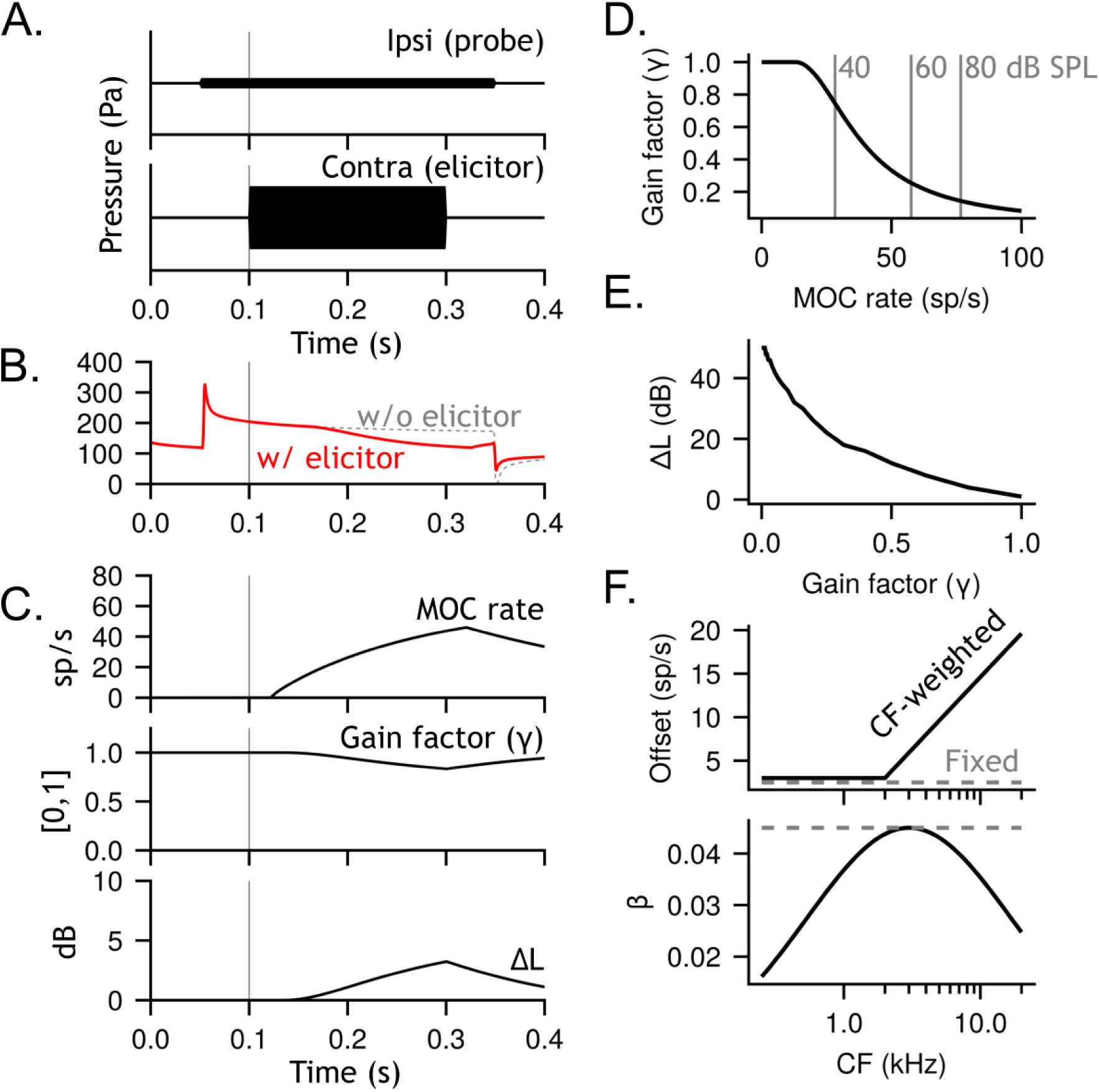
(A) Stimulus waveforms for the CAS paradigm, with a probe tone presented at 5 dB SPL (top) or an elicitor tone presented at 70 dB SPL (bottom). Note that vertical axes are not to scale. The vertical gray line here, and in B and C, indicates the onset of the elicitor. (B) Simulated ipsilateral AN response for an 8-kHz HSR fiber in response to the stimuli in A, either with the MOC pathway disabled (dashed gray) or the MOC pathway enabled (red). (C) Responses at intermediate model stages underlying MOC gain control in response to the stimuli in A. The top panel shows lowpass-filtered LSR rates from the 8-kHz contralateral CF, used as a proxy for WDR information in the ascending auditory pathway. The middle panel shows the resulting “gain factor” γ bottom panel shows the effective time-varying change in cochlear gain corresponding to each γ in the same channel based on the MOC IO nonlinearity and following the spatial-mixing step. The value in time, as inferred from midpoint shifts in on-CF RLFs (see Sec. IIC for details). (D) The IO nonlinearity relating MOC rate to gain factor at the 8-kHz CF. Note that this subfigure shows the single-channel gain factor before the spatial-mixing step; in practice, the mixing step substantially attenuated final gain factors, at least for pure-tone stimulation for which stimulation was highly tonotopically specific. (E) The relationship between gain factor and effective change in cochlear gain (ΔL), as mentioned in C. (F) Parameters in the MOC model as a function of CF, either for the version with the same parameters at all CFs (gray dashed) or CF-varying parameters (black). The IO “β”) on the bottom. threshold parameter (labeled “Offset”) is shown on the top and the IO slope parameter (labelled “β”) on the bottom.

#### 2. “Suppressor-level” functions

The threshold and magnitude of the model CAS reflex were characterized by simulating responses to ipsilateral pure-tone probes at a fixed sound level presented during ongoing contralateral elicitors at varying sound levels. Stimulus durations, ramp durations, and temporal configurations matched those used to simulate RLFs, as described above and depicted in Fig. 2. Typically, elicitor levels were varied between 10–70 dB SPL in steps of 2.5 dB SPL, while the probe was always presented at the midpoint of the ipsilateral RLF at CF (this choice matched that used in physiology and ensured that a contralateral elicitor’s effects were always visible in AN rates, which would not be the case if the probe were presented below threshold or above levels yielding rate saturation). Rate versus suppressor-level data were interpolated via linear interpolation as not all functions were well fit by a sigmoid. Subsequent analyses (e.g., identification of CAS threshold, see Sec. III) were always performed on the interpolated results.

#### 3. Tuning curves

Tuning curves involved simulating RLFs at probe frequencies centered logarithmically around CF, estimating the threshold for each RLF according to the method described above, and then smoothing the resulting threshold versus frequency data via locally estimated scatterplot smoothing (loess) using second-order polynomials with a relative span of 0.2. Tuning curves were only estimated for single-channel baseline models (i.e., without MOC gain control). In most cases, 71 probe frequencies ranging from three octaves below CF to three octaves above CF were tested, with levels spanning from 0 to 70 dB SPL in increments of 2.5 dB used to estimate tuning curves (in some cases, a higher maximum level was used when needed).

#### 4. CAS frequency sweeps

The effect of contralateral elicitor frequency on ipsilateral probe responses was assessed by simulating average-rate responses for a 100-ms probe presented at the midpoint of the ipsilateral RLF at the end of 1-s pure-tone contralateral elicitors at 65 dB SPL with varying elicitor frequencies. For visualizations of the resulting curve in comparison to physiological data, 21 tone frequencies ranging from three octaves below to three octaves above the ipsilateral CF were tested, and resulting probe-rate versus contralateral-elicitor-frequency data were smoothed using loess with a relative span of 0.25. For determination of “best suppressor frequency” (BSF), or the contralateral elicitor frequency that maximally reduced the probe rate response, contralateral elicitor frequencies were fixed at 41 values spanning logarithmically from 0.18–8 kHz (rather than being yoked to ipsilateral CF) and tested with 15 tested ipsilateral CF values spanning logarithmically from 0.5–16 kHz. The BSF for each ipsilateral CF was defined as the frequency that elicited the lowest probe rate on the smoothed data curve.

## III. RESULTS

### A. Simulating contralateral acoustic stimulation in the AN

We simulated the effects of CAS on AN responses via a “quasi-binaural” model that makes simplifying assumptions about CAS effects (see Sec. II for details; Fig 1D). When the MOC pathway was disabled, ipsilateral model fibers responded only to the ipsilateral pure-tone probe (Fig. 2A, top) and were unaffected by the contralateral pure-tone elicitor (Fig. 2A, bottom), demonstrating a characteristic onset response followed by adaptation to a quasi-steady state (Fig. 2B, dashed gray). In contrast, when the MOC pathway was enabled, the ipsilateral model fiber still responded directly only to the pure-tone probe; however, this response was modulated by MOC gain control arising from the contralateral elicitor (Fig. 2B, red). When the elicitor was gated on after the probe, the initial probe response was identical both with and without the MOC system enabled. With the MOC system enabled, changes in the probe response became visible after the 25-ms delay in the MOC pathway and after the sluggish MOC responses^3^ had accumulated (Fig. 2C, top). MOC responses had to surpass the threshold in the input-output (IO) nonlinearity mapping from MOC rate to gain-control factor to affect probe responses (Fig. 2D; see Sec. II for details). The resulting time course was an interplay between the lowpass filter in the MOC model and adaptation dynamics in the AN model itself. Qualitatively, this time course resembled the observed time course of CAS on AN responses (Warren and Liberman, 1989a).

### B. Contralateral acoustic stimulation thresholds

We began by simulating CAS thresholds as a function of CF and model parameters (Fig. 3). Warren and Liberman (1989a) define a unit’s CAS threshold as the elicitor level at which a probe-with-elicitor response can be distinguished from probe-alone response by eye. When measuring a CAS threshold, Warren and Liberman matched probe frequency to the unit’s CF and probe level to the unit’s RLF midpoint (Fig. 3A). Using the same approach, we operationally defined CAS threshold as the elicitor level yielding a 5% reduction in probe rate (Fig. 3B). Because stochastic aspects of the model (e.g., random spiking) were removed for the present simulations (see Sec. II), model CAS thresholds were a deterministic function of the model parameters.

**Figure 3.**
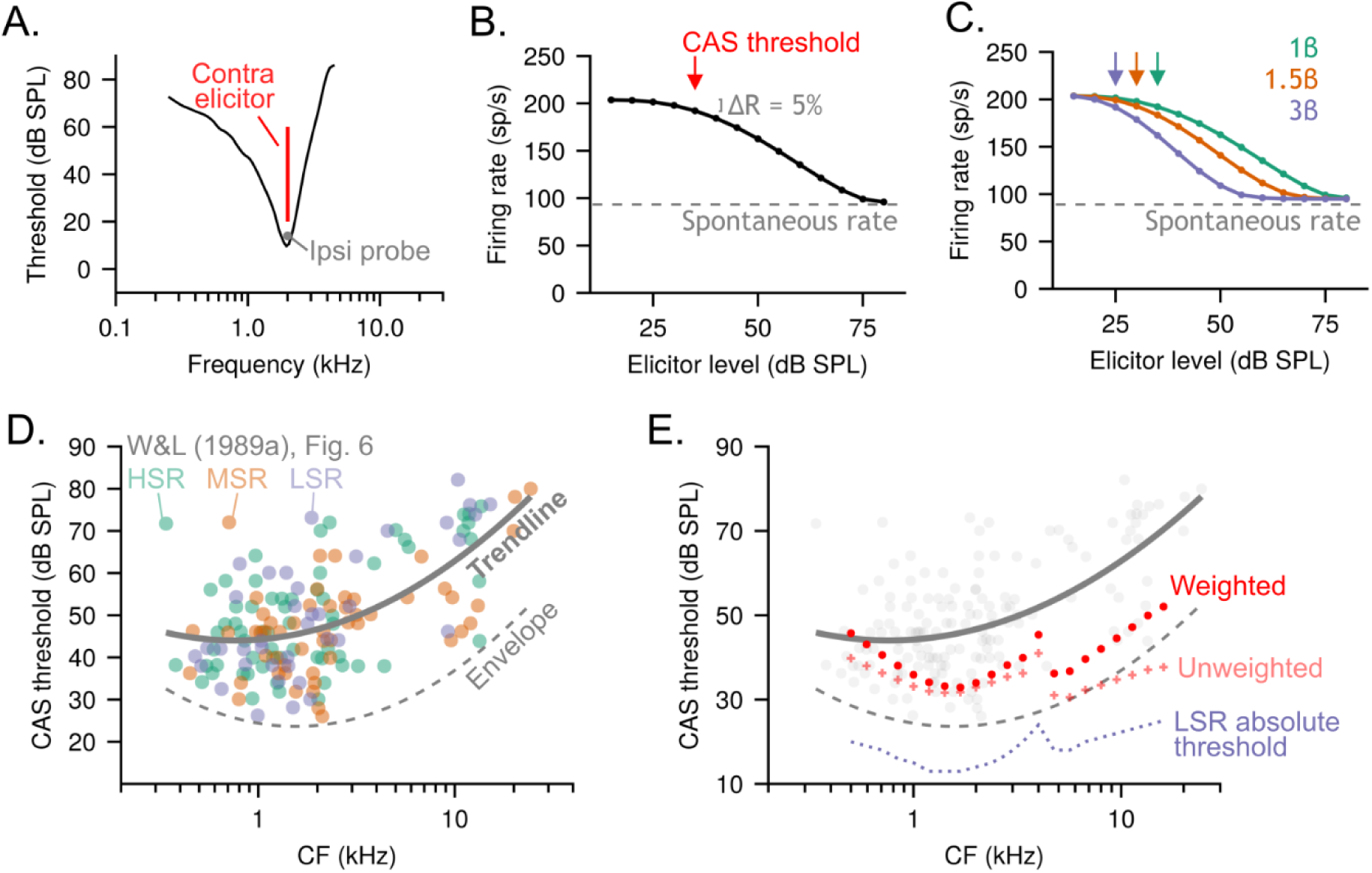
(A) Example iso-response tuning curve for a 2-kHz model HSR fiber (black line) in comparison to the frequency and level of a probe stimulus (gray marker) and frequency and level range of an elicitor stimulus (red line) used to determine the CAS threshold. (B) Corresponding firing rate versus elicitor level curve for the model neuron in A, with a red arrow indicating CAS threshold (elicitor level achieving 5% reduction in probe rate). (C) Effect of the MOC IO nonlinearity parameter, relative to the default value, on the curve shown in B. (D) CAS thresholds as function of CF from Warren and Liberman (1989a), Fig. 6. Colored markers indicate individual-fiber data; the solid gray line a trendline minimizing mean-squared-error across all fiber types; the dashed gray line a “lower envelope” of the data. (E) Data from A replotted in gray alongside model LSR thresholds as a function of CF (dashed purple line) and CAS thresholds for a model HSR fiber (solid red line).

Physiological data pooled across animals and units show a wide spread in CAS thresholds at a given CF (Warren and Liberman, 1989a). This spread reflects a combination of within-animal variance (e.g., differences within animal across units in threshold) and across-animal variance (e.g., differences in overall hearing sensitivity across animals). We did not seek to capture all of this variability with the model, and instead only to capture the general trends shared across animals and units. To this end, we first estimated a parametric lower “envelope” of the data (Fig. 3D, dashed gray line) and a parametric trendline through the data (Fig. 3D, solid gray line; see Sec. II for details). We reasoned that our model should (1) not produce responses that fall outside of the observed range of physiological data (i.e., outside the envelope), (2) display the general relationship between CAS threshold and CF suggested by both the envelope and the trendline, and (3) fall between the envelope and the trendline, consistent with the goal of representing a “best case scenario” in terms of overall sensitivity and strength of efferent responses. A similar reasoning was applied to other comparisons to physiological data presented below.

Empirical CAS thresholds are lowest for units with CFs near 1–2 kHz and comparatively higher for units with higher CFs (Warren and Liberman, 1989a; Fig 3D). When our MOC pathway had the same parameter values at all CFs (Fig. 2F, dashed gray), model CAS thresholds did not exhibit the correct trend with respect to CF or were too sensitive compared to the physiological data (Fig. 3E, pink). Model CAS thresholds were largely determined by a few key factors at a given CF: (1) the afferent LSR threshold (Fig. 3E, purple), (2) the properties of the LSR RLF, such as slope (Supp. Figs. 1A), (3) the amount of cochlear gain available (Supp. Fig. 1B), and (4) the parameters of the MOC IO nonlinearity. Model LSR thresholds varied with CF in a way that was consistent with physiological data (Liberman, 1978), reflecting primarily the effects of middle-ear filtering. As can be seen in Fig. 3E, the unweighted CAS thresholds generally mimicked the contours of the LSR thresholds with respect to CF, except that the gap between LSR thresholds and CAS thresholds was smaller at high CFs. This smaller gap at high CFs reflected the greater steepness of LSR RLFs at high CFs in our model (Supp. Fig. 1A) as well as the greater amount of cochlear gain available at high CFs (Supp. Fig. 1B). As a result of the slope difference, a unit increase in elicitor sound level a unit change in γ resulted in a larger effective reduction in cochlear gain at high than at low CFs. will resulted in a larger change in LSR rate at high than at low CFs. As a result of the gain difference, These differences combined to produce qualitatively different CAS thresholds (relative to LSR thresholds) at low and high CFs.

To address the discrepancy between how model and empirical CAS thresholds varied with CF, we heuristically adjusted both the IO threshold parameter and the slope parameter in the MOC IO nonlinearity as a function of CF (Fig. 2F, black). We used simple functional forms to vary IO threshold and slope across the CF range to qualitatively approximate a match to the physiological CAS data, rather than fitting the parameters with a nonlinear optimizer. These CF-varying parameters provided a better fit to the physiological CAS thresholds at high CFs (Fig. 3E, pink vs red). The fit to the data remained imperfect because the parameter values selected were a compromise intended not only to match the CAS thresholds but also other model responses (e.g., RLF shifts, see below).

Our final CAS thresholds exhibited a salient notch at CFs near 4 kHz, originating from the notch in the cat middle-ear filter transfer function (Rosowski, 1991) (Supp. Fig. 1A). It is challenging to compare this result directly to the data, as there is a relative sparsity of physiological data available at similar CFs; to avoid contaminating the results with the effects of acoustic crosstalk, the presence of these notches in the physiological experiments often precluded the presentation of precursors at high enough sound levels to evoke a reflex (Warren and Liberman, 1989a). For other species middle-ear filter transfer functions, this notch in the CAS thresholds would be absent.

### C. Rate-level function changes

Next, we evalauted how stimulation of the MOC system via contralateral sound affected afferent RLFs in the model, and whether these effects were consistent with available physiological data (Warren and Liberman, 1989a, 1989b). RLFs were described using several features: (1) the rate range was defined as the firing rates spanning 10 to 90% of the extrema firing rates (Fig. 4B, vertical bar); (2) the dynamic range was defined as the span of the sound levels corresponding to the rate range (Fig. 4B, horizontal bar); (3) the threshold was defined as the level achieving the minimum of the rate range (Fig. 4B, arrow marked “TH”); and (4) the midpoint was defined as the level achieving the middle of the rate range (Fig. 4B, arrow marked “MP”).

**Figure 4.**
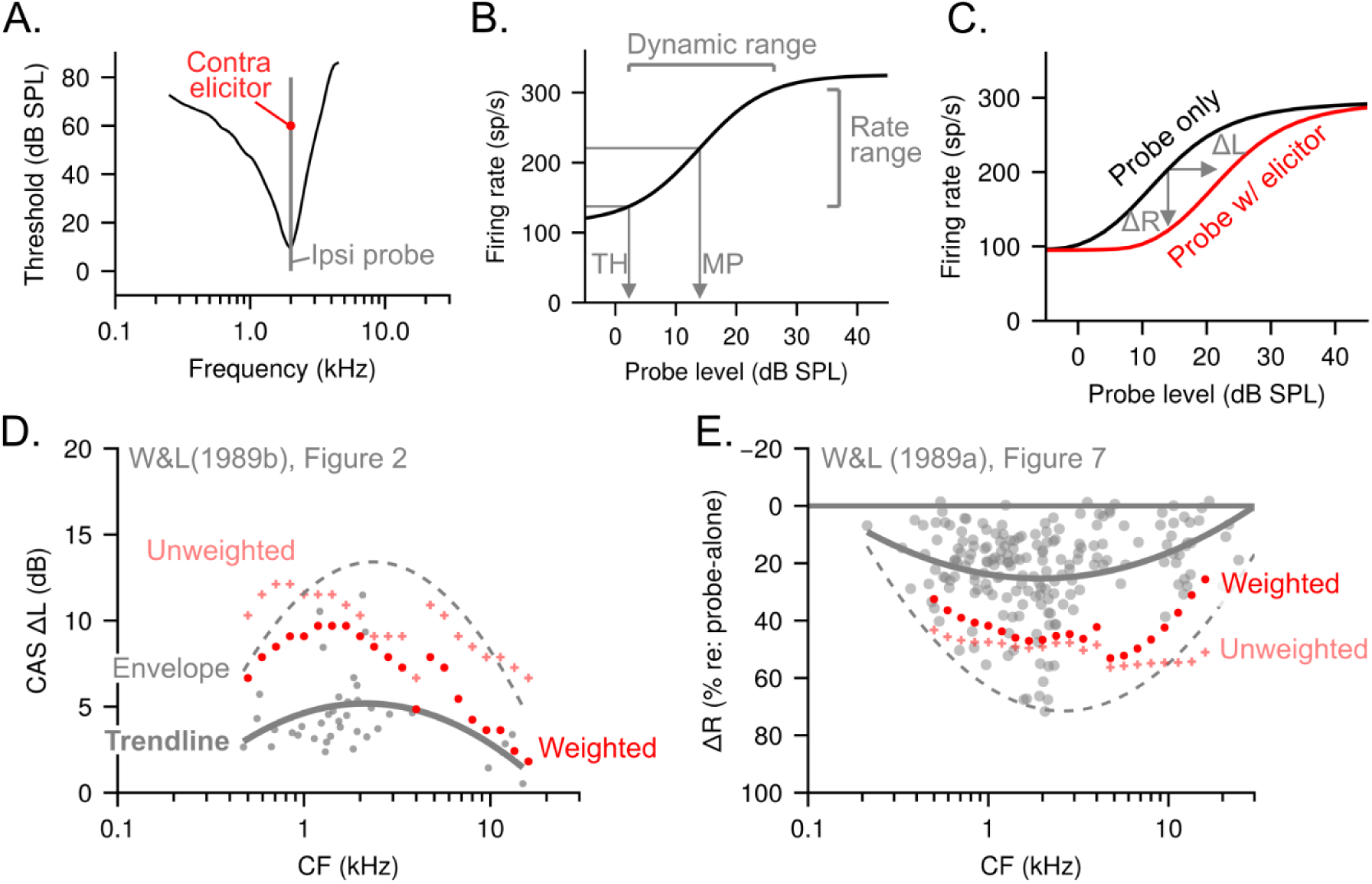
(A) Example tuning curve for a 2-kHz model HSR fiber (black line) in comparison to the frequency and level of a probe stimulus (gray line) and frequency and level range of an elicitor stimulus (red marker) used to determine ΔL and ΔR from an RLF simulation. (B) Corresponding probe-alone RLF for the model neuron in A. Gray arrows indicate rate threshold (TH) and RLF midpoint (MP), while gray brackets indicate the 10–90% rate range and the dynamic range, as defined in Sec II. (C) Probe-alone RLF from B (black line) versus a corresponding probe-with-elicitor RLF (red line). ΔL and ΔR are marked with horizontal and vertical gray arrows, respectively. (D) ΔL for elicitors in the range of 60–85 dB SPL from Figure 2 of Warren and Liberman (1989b). Markers indicate individual-fiber ΔLs. The solid gray line is a trendline minimizing mean-squared-error across all fiber types. The dashed gray line is an “upper envelope” of the data. Markers and trendlines in red show simulated ΔLs. (E) ΔR for elicitors in the range of 60–70 dB SPL from Figure 7 of Warren and Liberman (1989a). Markers indicate individual-fiber ΔRs. The solid gray line is a trendline minimizing mean-squared-error across all fiber types. The dashed gray line is an “upper envelope” of the data. Markers and trendlines in red show simulated ΔRs.

Our baseline model (i.e., the model with MOC disabled) inherited the good correspondence of simulated RLFs to empirical RLFs present in the earlier versions of the peripheral model (Zilany et al., 2009, 2014). Specifically, for an HSR fiber, simulated RLFs have sensitive thresholds, limited dynamic ranges, and strong rate saturation (Fig. 4B). When the model MOC system was enabled, adding a contralateral elicitor at the appropriate frequency and sound level shifted the RLF rightward due to efferent gain control (Fig. 4C, red versus black lines). This basic pattern of a rightward shift in the RLF with little change in the slope was consistent with both CAS-evoked and electrically evoked MOC responses (Guinan and Gifford, 1988a; Warren and Liberman, 1989b). When comparing two RLFs, we summarize the changes via the shift in midpoint (Fig. 2C, arrow marked “ΔL”) and the reduction in rate for a probe at the RLF midpoint (Fig. 2C, arrow marked “ΔR”).

With the CF-dependent IO threshold and slope applied to reduce the strength of MOC effects at low and high CFs (Fig. 2F), both ΔL (Fig. 4D) and ΔR (Fig. 4E) simulations were satisfactory. For the ΔL simulations, simulated values fell within the range of observed data and followed the overall trend seen in the data, with the largest effects near CFs of 2 kHz and weaker effects at more distant CFs (Warren and Liberman, 1989a, 1989b). For the ΔR simulations, the results showed more deviation from the physiological data, with the largest effects at CFs near 4-5 kHz, instead of 2 kHz. The ΔR results peaked at this higher CF because in the baseline model HSR RLFs become very steep at high CFs, so that a unit change in gain results in a larger change in rate for higher versus lower CFs (Supp. Fig. 1A).

Importantly, and consistent with the physiological data, these effects were observed without producing changes in either the dynamic range or the rate range of RLFs. Averaged across CF, dynamic range without a precursor was 20.0 dB, while dynamic range with a precursor was 20.2 dB. Likewise, averaged across CF, rate range without a precursor was 221 Hz, while rate range with a precursor was 227 Hz. That is, CAS resulted in pure rightward shifts of RLFs, and did not substantially change RLF slope or dynamic range.

### D. Off-CF elicitors and other multiband effects

Next, we investigated the consequences of the multichannel design of our efferent model. As discussed in Sec. II, the present model differed from other filterbank models of auditory processing in that it simulated multiple channels in a single fused time loop. This design allowed responses from multiple peripheral channels to be combined in the feedforward direction to influence responses in other channels of subsequent stages, or for responses from individual model MOC channels to influence responses in multiple peripheral channels in a feedback direction (i.e., multichannel gain control). In our model, each channel produced a single gain control factor γ based on the mapping between lowpass-filtered LSR rates rate and gain-control factors. Single-channel gain-control factors were then spatially mixed along the (log) tonotopic axis using a Gaussian weighting function before determining cochlear gain at subsequent time steps. This multichannel formulation of MOC gain control was motivated by both anatomical and physiological evidence. Anatomically, individual MOC neurons innervate OHCs in the cochlea over sometimes fairly wide tonotopic spans (Brown, 2014; Liberman and Brown, 1986). This observation suggests that, while the overall MOCR circuit is tonotopic and constituent neurons in the circuit are all sharply tuned, MOC gain control may nevertheless be “blurred” in terms of the frequency specificity of its effects in the cochlea.

Physiologically, contralateral elicitor tones can elicit significant suppression of ipsilateral probe rates over a wide frequency range in individual afferent neurons (Warren and Liberman, 1989b, their Fig. 6; reproduced here in Fig. 5B). Similar evidence of broad frequency effects in MOC gain control are seen in otoacoustic emissions studies (Lilaonitkul and Guinan, 2009).

**Figure 5.**
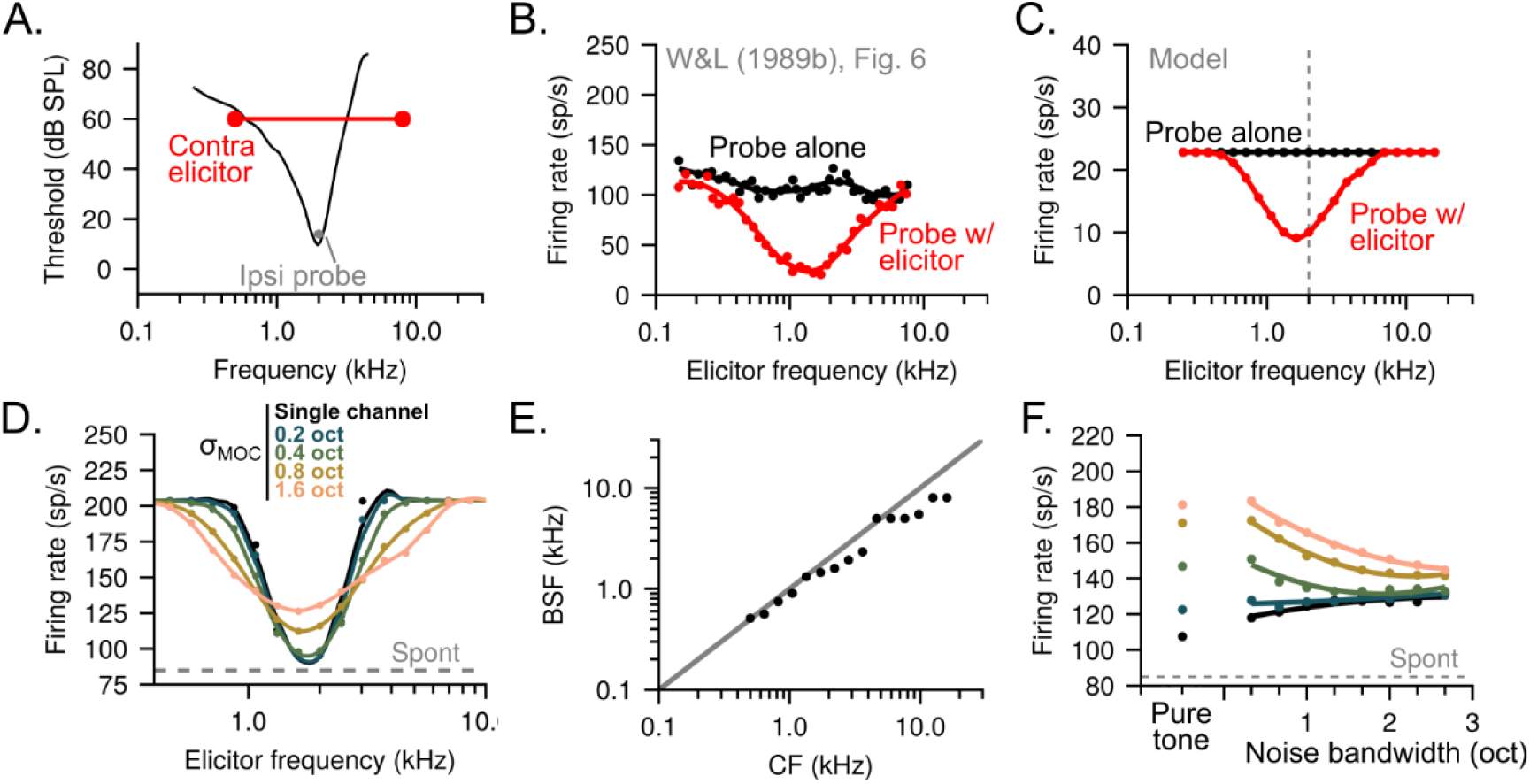
(A) Example tuning curve for a 2-kHz model HSR fiber (black line) in comparison to the frequency and level of a probe stimulus (gray marker) and level and frequency range of an elicitor stimulus (red line and markers) used to characterize the tuning characteristics of MOC gain control in the model. (B) Reproduction of data from Warren and Liberman (1989b), Figure 6. (C) Simulation of results from B using a 2-kHz LSR fiber. Probe level matched the midpoint of the 2-kHz RLF while elicitor level was fixed at 70 dB SPL. (D) Effect of the MOC mixing bandwidth, expressed in units of standard deviation in octaves, on result shown in C but with an HSR fiber, with different values indicated by different colors. (E) Scatter plot of the best suppressor frequency (BSF) vs CF for a range of CFs, given a 65 dB SPL pure-tone suppressor. (F) Effect of widening th bandwidth of a bandlimited noise suppressor on the result shown in C. Colors are as in D.

We simulated responses to contralateral pure-tone elicitors with frequencies varied over a wide range and compared the results to physiology. With a Gaussian mixing weight function with a standard deviation of 0.9 octaves, we found that our model exhibited rate reductions due to CAS in a 2-kHz LSR fiber over a frequency range comparable to the example neuron from Warren and Liberman (1989b). Specifically, significant rate reductions could be seen at least one octave below and above CF (Fig. 5C). When the spatial mixing of gain factors was eliminated or restricted to a narrower tonotopic range, CAS-induced probe-rate reduction extended over too narrow a frequency range, largely determined by the bandwidth of LSR afferent tuning, which drives the MOCR in the model (Fig. 5D). Note that data in Fig. 5B and the simulations in Figs. 5C and 5D had different absolute rates. Fig. 5B was recorded from an LSR neuron and Fig. 5C simulated from an LSR neuron, but their rates are quite different because our model LSR fiber at 2 kHz has a relatively low driven rate (Supplementary Figure 1), while Fig. 5D was simulated from an HSR neuron. Nevertheless, because probes were always presented at the midpoint of each unit’s RLF, the effects on probe rate were qualitatively similar across the three cases, shifting probe rates between midpoint rate and spontaneous rate based on cochlear gain reduction.

The CAS elicitor frequency that yielded the greatest reduction in ipsilateral rate, or best suppressor frequency (BSF), was closely related to ipsilateral CF with a few stereotyped deviations from a one-to-one trendline (Fig. 5E). First, BSF was sometimes slightly lower than CF, which may arise from asymmetries in the peripheral filters and the shift of best frequency vs characteristic frequency at higher sound levels. A similar pattern was observed by Warren and Liberman (1989b). Second, there was a noticeable deviation from a linear relationship between BSF and CF at CFs near 4 kHz due to the notch in the cat middle-ear filter used in the model. For example, an elicitor presented at 2 kHz elicited a larger overall response in the LSR population than an elicitor presented at 4 kHz because of the notch. If the mixing of gain factors across CF were sufficiently broad, then an HSR fiber tuned to 4 kHz might experience more gain control under the 2 kHz than the 4 kHz elicitor. As a result, its BSF would be noticeably lower than CF.

The multichannel design of our model leads to interesting predictions for non-tonal elicitors. In a simple same-channel MOCR model, wherein each channel exclusively regulates its own gain, increases in elicitor energy that fall outside the passband of the channel’s peripheral filter should have little effect on gain control. In other words, in such models, MOC gain control should have an “elicitor critical band” that reflects the tuning bandwidth of the peripheral filter driving it. In contrast, in the present multichannel design, elicitor energy outside the passband of a given channel’s peripheral filter could still influence gain control in that channel. To illustrate this, we simulated probe responses in an HSR fiber with a 2-kHz CF with either a pure-tone elicitor or a noise elicitor of varying bandwidths centered on 2 kHz (Fig. 5F). The noise elicitors had a constant spectrum level of 20 dB SPL, and the pure tone elicitor had a level matching the overall level of the 1/3-octave noise elicitor (49 dB SPL). In the single-channel MOC model, gain control was relatively strong for the pure-tone elicitor, because the lack of a spatial-mixing step in calculating gain-control factors resulted in less gain control “leaking” to other channels (Fig. 5F, black marker). When the pure-tone elicitor was replaced with a level-matched 1/3-octave noise elicitor, slightly less gain control was elicited and probe rates were slightly higher. As the bandwidth of the noise elicitor was increased, even less gain control was elicited and probe rates continued to rise (Fig. 5F, black line). This pattern may have arisen from peripheral suppression in the model. The added energy in the wider elicitors fell outside the passband of the sharply tuned signal-path filter in the model, and thus could not contribute to MOC gain control in the single-channel model. However, some of that energy fell within the passband of the more broadly tuned control-path filter that regulates the signal-path filter gain to emulate two-tone suppression, reducing its sensitivity and thus the magnitude of the afferent signal driving the MOCR. In contrast, in multiband models, wherein gain control factors were spatially mixed over the tonotopic range, pure-tone elicitors resulted in the highest probe rates (least gain control), whereas increasingly noise elicitors resulted in lower probe rates (more gain control) and probe rates dropped systematically with increasing elicitor bandwidth (Fig. 5F, colored lines). This result is consistent with observations that broadband elicitors are more effective at eliciting the MOCR than narrowband elicitors (Lilaonitkul and Guinan, 2009) and suggests that an important ingredient in achieving this result may be modeling the “wideband” nature of MOC gain control.

## IV. DISCUSSION

We presented a new model of the early auditory system that simulates efferent feedback from the brainstem to the OHCs via the MOC pathway. The model exhibited the hallmarks of MOC gain control elicited by CAS as observed via single-unit AN physiology (Warren and Liberman, 1989a, 1989b). That is, reductions in cochlear gain due to MOC gain control manifested as elevations in afferent thresholds and rightward shifts of afferent RLFs at CF. MOC gain control could first be observed in the model for contralateral elicitors in the range of 35–50 dB SPL (Fig. 3). Elicitors near 65 dB SPL yielded rightward shifts in RLFs of up to 10 dB and reductions in response rate on the order of 40% for HSR fibers (Fig. 4). MOC effects were strongest for CFs near 2 kHz and weaker elsewhere, particularly at high CFs (Figs. 3 and 4). Significant reductions in probe rate at CF could be observed over a wide range of contralateral elicitor frequencies relative to CF (Fig. 5).

The present MOC model has major limitations that are worth addressing. One key limitation is that the model is phenomenological, in that it seeks only to reproduce the relationships observed in data and not to describe the biophysical processes that realize those relationships (Osses Vecchi et al., 2022). As such, detailed representations of intermediate stages are elided in favor of a simple abstraction of the underlying computations; lowpass-filtered LSR rates serve as a proxy for suprathreshold level information presumed to drive MOC activity, and the operating range and behavior of the MOCR are mimicked with filters and static nonlinearities. Future models will need to synthesize available data on the MOCR circuit to model the anatomy and physiology of its constituent components in more detail. Level information needed to govern the MOCR is clearly available to MOC neurons, insofar as MOC neurons (above their thresholds near 40 dB SPL) exhibit dynamic ranges for pure-tone stimulation that readily exceed 40 dB (Liberman and Brown, 1986).

Several sources of evidence implicate cells in the posteroventral cochlear nucleus, planar multipolar/T-stellate neurons, as a primary source of excitatory input to the MOC (Benson and Brown, 2006; Brown et al., 2013; Darrow et al., 2012; De Venecia et al., 2005; Liberman and Brown, 1986; Romero and Trussell, 2021), although Carney (2018) argues that the limited dynamic range and threshold variability of T-stellates would not yield a robust code for level over the sound levels where the MOCR is active. Other sources of level information to the MOC may include neurons in the “small cell cap” of the VCN that are thought to receive input primarily from medium-and low-SR AN fibers (Hockley et al., 2022; Ryugo, 2008; Ye et al., 2000). Future models that incorporate more details of these pathways are likely to fare better in capturing the nuances of MOC gain control.

The present MOC model inherits the assumptions of the Zilany-Bruce-Carney series of auditory peripheral models (Bruce et al., 2003; Zilany et al., 2009, 2014), and thus also its strengths and weaknesses. Perhaps the most important of these assumptions is that the auditory system can be adequately modeled as a parallel filterbank, as opposed to explicitly simulating the spatially coupled nature of cochlear mechanics (Altoè et al., 2014; Verhulst et al., 2018). As emerging evidence emphasizes the importance of such aspects of cochlear mechanics (Cooper et al., 2018; Kondylidis et al., 2025), it will be important to critically scrutinize filterbank approaches for simulating peripheral auditory processing. It is interesting to wonder—although at this point purely speculatively—how the notion of a spatially distributed cochlear amplifier might relate to the spatially broad innervation of OHCs by MOC fibers (Brown, 2014). Another key limitation of the 2014 version of the Zilany-Bruce-Carney model (which was partially addressed in the model of Bruce et al. [2018]), is that different fiber types (e.g., HSR vs LSR) have fixed, stereotyped thresholds and dynamic ranges. This simplification stands in contrast to the within-group diversity of threshold and dynamic ranges seen in AN fibers and limits the ability of the model to account for corresponding variation in CAS effects. The present model was adapted from the 2014 model rather than the 2018 model because changes to the 2018 model yielded improvements in the statistics of spike-train outputs at the cost of the accuracy of its instantaneous-rate outputs. The current model relies on the instantaneous-rate outputs to provide a continuous and non-stochastic control signal to govern the MOCR. A future model could reconcile these differences to provide usable MOC control signals while also benefitting from the improvements of the 2018 model vis-à-vis AN fiber threshold and spontaneous rate diversity.

The present MOC model assumes that the MOC system can be understood as an automatic gain control system where increases in stimulus energy—heuristically inferred from the activity of non-saturating LSR AN fibers—result in decreases in cochlear gain. This assumption is in keeping with classical theories of the MOC system, but we emphasize here that this is an incorrect assumption. In reality, MOC neurons do not respond to sound as simple level estimators. For example, MOC neurons have been shown to exhibit rate tuning to amplitude-modulation frequency (Gummer et al., 1988), possibly originating from strong descending inputs from inferior colliculus (Farhadi et al., 2023; Romero and Trussell, 2021, 2022). Nevertheless, our hope is that the present simplification of the problem is a useful first step in the development of more sophisticated and realistic models in the future. Along these lines, we reasoned that, because the stimuli used here (pure tones and Gaussian noise) are fairly simple, MOC neurons’ sensitivities to features of sound other than energy would play a negligible role in the overall results. Our lab has previously developed a different auditory model with MOC gain control that simulates the potential contribution of descending input from modulation-sensitive inferior-colliculus neurons to the MOC (Farhadi et al., 2023). Presently, these two efferent models can be understood as different leaves of a broader branch of models, each seeking to address different questions about the consequences of MOC gain control on peripheral coding based on different sources of data. Future models will have to reconcile these two modeling approaches as more data on MOC anatomy and physiology becomes available, particularly regarding the role of inputs to MOC beyond those in the classic brainstem-level three-synapse MOC-reflex circuit (Romero and Trussell, 2022).

Future versions of the model will need to more accurately simulate the binaural aspects of the MOC system. As summarized previously, gain in a given cochlea can be affected by sound presented to either ear via different pathways within the MOC system. In a CAS paradigm, MOC effects measured in the probe ear should be driven almost exclusively by that ear’s contralateral reflex (driven by the elicitor), as the probe is typically presented at sound levels that would not be expected to drive MOC neurons in the ipsilateral reflex (< 40 dB SPL). The case in the elicitor ear is different; the elicitor ear’s contralateral reflex (driven by the probe) should be weak, as the probe is usually at a relatively low sound level, but the elicitor ear’s ipsilateral reflex (driven by the elicitor) almost certainly influences the response to the elicitor itself and thus the overall effect of CAS. Qualitatively, we found that the effects of the elicitor ear’s ipsilateral reflex were nonnegligible for high elicitor levels (result not shown). The present model was designed to simulate CAS data, but CAS was simulated by approximation; the contralateral (elicitor) ear was simulated first in time and then its outputs were used to govern cochlear gain in the ipsilateral (probe) ear, precluding simulation of full dynamic interactions across the two ears. In brief, this strategy effectively simulated the contributions of both the elicitor ear’s ipsilateral reflex and the probe ear’s contralateral reflex. However, it did assume that the two reflexes are identical (e.g., in threshold, magnitude, temporal dynamics), whereas most evidence suggests important differences between the two (Guinan, 2014). This partial framework for simulating the MOCR will need to be replaced with a true binaural framework that properly models the differences between the ipsilateral and contralateral reflex pathways if the model is to be used to simulate the simultaneous contributions of the two reflexes in complex binaural sounds.

An intriguing dimension of future work is how to translate the present model, which is based on physiological data from cat, to humans. There is substantial variation in the physiology and anatomy of the MOC system across mammalian animal-model species (Lopez-Poveda, 2018; Warr, 1992), including in: number of MOC neurons (Warr, 1992), the sharpness of their tuning (Brown, 1989; Liberman and Brown, 1986), the relative balance of crossed and uncrossed pathways (Warr, 1992), and the frequencies (or CFs) exhibiting the greatest MOC effects (Guinan and Gifford, 1988a; Guinan and Stankovic, 1996; Lilaonitkul and Guinan, 2012; Warren and Liberman, 1989a). It remains to be seen how well a model calibrated to reproduce data from cat will generalize to other mammalian species. The influence of anesthesia on the MOC system is another related issue. Most measurements of MOC effects in non-human animals are made under anesthesia, and anesthesia is known to suppress the MOCR (Chambers et al., 2012). The present model sought to reproduce data from cats anesthetized with diallyl barbiturate in urethane (Warren and Liberman, 1989a, b); how well such data generalize to awake animals or animals under different forms of anesthesia is largely unknown, and will be an important area for future research.

Psychophysicists have long wondered whether MOC gain control can explain certain “temporal effects” in auditory perception, such as auditory overshoot (Jennings et al., 2011; Roverud and Strickland, 2015; Strickland, 2004). Study of the auditory afferent system, and its links with perception, has been facilitated by readily available and detailed computational models of the auditory periphery and ascending auditory pathway (e.g., recently, Bruce et al., 2018; Verhulst et al., 2018). We hope that the present auditory efferent model will provide similar assistance in the quest to link auditory behavior and auditory efferent physiology.

## SUPPLEMENTARY MATERIAL

See supplementary material at [URL will be inserted by AIP] depicting input-output profiles for relevant model stages versus CF.

## ACKNOWLEDGMENTS

This work was supported by NIH NIDCD R01-DC010813 (L.H.C.), NIH NIDCD F32-DC022143 (D.R.G.), and NIH NIDCD F32-DC022782 (A.F.). We would also like to thank the many individuals who attended the workshops on auditory efferent systems at the Hanse-Wissenschaftskolleg and provided invaluable feedback on preliminary versions of this work.

## DATA AVAILABILITY

Data and code used in this publication are available in several places. First, an archival version of the data and code is available on OSF at [[link]]. Second, the current version of the model code, along with full documentation and tools to report bugs and issues, is available in a GitHub repository: [[link]]. Users are not intended to interface directly with the C code available in this GitHub repository. Instead, third, we provide user-friendly “wrapper” functions in both MATLAB and Julia for community use [[cite]].

## Footnotes

1 Here we describe the general case, but this parameter varies with CF.

2 Throughout, we use the word “ipsilateral” to refer to the simulated ear responding to the probe stimulus, that is, where the recording electrode would be located in a physiological experiment. Likewise, we use the word “contralateral” to refer to the simulated ear responding to the elicitor stimulus, which would be on the opposite side of the head from the recording electrode in a physiological experiment.

3 As a convenience, we refer to the lowpass-filtered LSR rates as MOC rates or MOC responses.

## REFERENCES

1. Almishaal, A., Bidelman, G. M., and Jennings, S. G. (2017). “Notched-noise precursors improve detection of low-frequency amplitude modulation,” The Journal of the Acoustical Society of America, 141, 324–333. doi:10.1121/1.4973912

2. Altoè, A., Pulkki, V., and Verhulst, S. (2014). “Transmission line cochlear models: Improved accuracy and efficiency,” The Journal of the Acoustical Society of America, 136, EL302–EL308. doi:10.1121/1.4896416

3. Bachman, J. L., Kitcher, S. R., Vattino, L. G., Beaulac, H. J., Chaves, M. G., Hernandez Rivera, I., Katz, E., et al. (2025). “GABAergic synapses between auditory efferent neurons and type II spiral ganglion afferent neurons in the mouse cochlea,” Proc. Natl. Acad. Sci. U.S.A., 122, e2409921122. doi:10.1073/pnas.2409921122

4. Backus, B. C., and Guinan, J. J. (2006). “Time-course of the human medial olivocochlear reflex,” The Journal of the Acoustical Society of America, 119, 2889–2904. doi:10.1121/1.2169918

5. Benson, T. E., and Brown, M. C. (2006). “Ultrastructure of synaptic input to medial olivocochlear neurons,” J of Comparative Neurology, 499, 244–257. doi:10.1002/cne.21118

6. Brown, M. C. (1989). “Morphology and response properties of single olivocochlear fibers in the guinea pig,” Hearing Research, 40, 93–110. doi:10.1016/0378-5955(89)90103-2

7. Brown, M. C. (2001). “Response Adaptation of Medial Olivocochlear Neurons Is Minimal,” Journal of Neurophysiology, 86, 2381–2392. doi:10.1152/jn.2001.86.5.2381

8. Brown, M. C. (2011). “Anatomy of Olivocochlear Neurons,” In D. K. Ryugo, R. R. Fay, and A. N. Popper (Eds.), Auditory and Vestibular Efferents, Springer Handbook of Auditory Research, Springer, pp. 17–37.

9. Brown, M. C. (2014). “Single-unit labeling of medial olivocochlear neurons: the cochlear frequency map for efferent axons,” Journal of Neurophysiology, 111, 2177–2186. doi:10.1152/jn.00045.2014

10. Brown, M. C., De Venecia, R. K., and Guinan, J. J. (2003). “Responses of medial olivocochlear neurons: Specifying the central pathways of the medial olivocochlear reflex,” Exp Brain Res, 153, 491–498. doi:10.1007/s00221-003-1679-y

11. Brown, M. C., Mukerji, S., Drottar, M., Windsor, A. M., and Lee, D. J. (2013). “Identification of Inputs to Olivocochlear Neurons Using Transneuronal Labeling with Pseudorabies Virus (PRV),” JARO, 14, 703–717. doi:10.1007/s10162-013-0400-5

12. Brown, M. C., and Nuttall, A. L. (1984). “Efferent control of cochlear inner hair cell responses in the guinea-pig.,” The Journal of Physiology, 354, 625–646. doi:10.1113/jphysiol.1984.sp015396

13. Brown, M. C., Nuttall, A. L., and Masta, R. I. (1983). “Intracellular Recordings from Cochlear Inner Hair Cells: Effects of Stimulation of the Crossed Olivocochlear Efferents,” Science, 222, 69–72. doi:10.1126/science.6623058

14. Bruce, I. C., Erfani, Y., and Zilany, M. S. A. (2018). “A phenomenological model of the synapse between the inner hair cell and auditory nerve: Implications of limited neurotransmitter release sites,” Hearing Research, 360, 40–54. doi:10.1016/j.heares.2017.12.016

15. Bruce, I. C., Sachs, M. B., and Young, E. D. (2003). “An auditory-periphery model of the effects of acoustic trauma on auditory nerve responses,” The Journal of the Acoustical Society of America, 113, 369–388. doi:10.1121/1.1519544

16. Carney, L. H. (2018). “Supra-threshold hearing and fluctuation profiles: Implications for sensorineural and hidden hearing loss,” Journal of the Association for Research in Otolaryngology, 19, 331–352. doi:10.1007/s10162-018-0669-5

17. Chambers, A. R., Hancock, K. E., Maison, S. F., Liberman, M. C., and Polley, D. B. (2012). “Sound-Evoked Olivocochlear Activation in Unanesthetized Mice,” JARO, 13, 209–217. doi:10.1007/s10162-011-0306-z

18. Cooper, N. P., and Guinan, J. J. (2006). “Efferent-mediated control of basilar membrane motion,” The Journal of Physiology, 576, 49–54. doi:10.1113/jphysiol.2006.114991

19. Cooper, N. P., Vavakou, A., and Van Der Heijden, M. (2018). “Vibration hotspots reveal longitudinal funneling of sound-evoked motion in the mammalian cochlea,” Nat Commun, 9, 3054. doi:10.1038/s41467-018-05483-z

20. Darrow, K. N., Benson, T. E., and Brown, M. C. (2012). “Planar multipolar cells in the cochlear nucleus project to medial olivocochlear neurons in mouse,” J of Comparative Neurology, 520, 1365–1375. doi:10.1002/cne.22797

21. De Venecia, R. K., Liberman, M. C., Guinan, J. J., and Brown, M. C. (2005). “Medial olivocochlear reflex interneurons are located in the posteroventral cochlear nucleus: A kainic acid lesion study in guinea pigs,” J of Comparative Neurology, 487, 345–360. doi:10.1002/cne.20550

22. Dolan, D. F., Guo, M. H., and Nuttall, A. L. (1997). “Frequency-dependent enhancement of basilar membrane velocity during olivocochlear bundle stimulation,” The Journal of the Acoustical Society of America, 102, 3587–3596. doi:10.1121/1.421008

23. Doucet, J. R., Rose, L., and Ryugo, D. K. (2002). “The cellular origin of corticofugal projections to the superior olivary complex in the rat,” Brain Research, 925, 28–41. doi:10.1016/S0006-8993(01)03248-6

24. Elgueda, D., Delano, P. H., and Robles, L. (2011). “Effects of Electrical Stimulation of Olivocochlear Fibers in Cochlear Potentials in the Chinchilla,” JARO, 12, 317–327. doi:10.1007/s10162-011-0260-9

25. Farhadi, A., Jennings, S. G., Strickland, E. A., and Carney, L. H. (2023). “Subcortical auditory model including efferent dynamic gain control with inputs from cochlear nucleus and inferior colliculus,” The Journal of the Acoustical Society of America, 154, 3644–3659. doi:10.1121/10.0022578

26. Faubion, S. L., Park, R. K., Lichtenhan, J. T., and Jennings, S. G. (2024). “Effects of contralateral noise on envelope-following responses, auditory-nerve compound action potentials, and otoacoustic emissions measured simultaneously,” The Journal of the Acoustical Society of America, 155, 1813–1824. doi:10.1121/10.0025137

27. Grange, J., Zhang, M., and Culling, J. (2022). “The Role of Efferent Reflexes in the Efficient Encoding of Speech by the Auditory Nerve,” J. Neurosci., 42, 6907–6916. doi:10.1523/jneurosci.2220-21.2022

28. Guest, D. R., and Carney, L. H. (2024). “A fast and accurate approximation of power-law adaptation for auditory computational models,” The Journal of the Acoustical Society of America, 156, 3954–3957. doi:10.1121/10.0034457

29. Guinan, J. J. (2006). “Olivocochlear Efferents: Anatomy, Physiology, Function, and the Measurement of Efferent Effects in Humans,” Ear and Hearing, 27, 589–607. doi:10.1097/01.aud.0000240507.83072.e7

30. Guinan, J. J. (2014). “Olivocochlear efferent function: Issues regarding methods and the interpretation of results,” Frontiers in Systems Neuroscience, 8, nil. doi:10.3389/fnsys.2014.00142

31. Guinan, J. J. (2018). “Olivocochlear efferents: Their action, effects, measurement and uses, and the impact of the new conception of cochlear mechanical responses,” Hearing Research, 362, 38–47. doi:10.1016/j.heares.2017.12.012

32. Guinan, J. J., and Gifford, M. L. (1988a). “Effects of electrical stimulation of efferent olivocochlear neurons on cat auditory-nerve fibers. I. Rate-level functions,” Hearing Research, 33, 97–113. doi:10.1016/0378-5955(88)90023-8

33. Guinan, J. J., and Gifford, M. L. (1988b). “Effects of electrical stimulation of efferent olivocochlear neurons on cat auditory-nerve fibers. II. Spontaneous rate,” Hearing Research, 33, 115–127. doi:10.1016/0378-5955(88)90024-X

34. Guinan, J. J., and Gifford, M. L. (1988c). “Effects of electrical stimulation of efferent olivocochlear neurons on cat auditory-nerve fibers. III. Tuning curves and thresholds at CF,” Hearing Research, 37, 29–45. doi:10.1016/0378-5955(88)90075-5

35. Guinan, J. J., and Stankovic, K. M. (1996). “Medial efferent inhibition produces the largest equivalent attenuations at moderate to high sound levels in cat auditory-nerve fibers,” The Journal of the Acoustical Society of America, 100, 1680–1690. doi:10.1121/1.416066

36. Gummer, M., Yates, G. K., and Johnstone, B. M. (1988). “Modulation transfer function of efferent neurones in the guinea pig cochlea,” Hearing Research, 36, 41–51. doi:10.1016/0378-5955(88)90136-0

37. Hockley, A., Wu, C., and Shore, S. E. (2022). “Olivocochlear projections contribute to superior intensity coding in cochlear nucleus small cells,” The Journal of Physiology, 600, 61–73. doi:10.1113/jp282262

38. Hua, Y., Ding, X., Wang, H., Wang, F., Lu, Y., Neef, J., Gao, Y., et al. (2021). “Electron Microscopic Reconstruction of Neural Circuitry in the Cochlea,” Cell Reports, 34, 108551. doi:10.1016/j.celrep.2020.108551

39. Jennings, S. G. (2021). “The role of the medial olivocochlear reflex in psychophysical masking and intensity resolution in humans: a review,” Journal of Neurophysiology, 125, 2279– 2308. doi:10.1152/jn.00672.2020

40. Jennings, S. G., Chen, J., Fultz, S. E., Ahlstrom, J. B., and Dubno, J. R. (2018). “Amplitude modulation detection with a short-duration carrier: Effects of a precursor and hearing loss,” The Journal of the Acoustical Society of America, 143, 2232–2243. doi:10.1121/1.5031122

41. Jennings, S. G., Heinz, M. G., and Strickland, E. A. (2011). “Evaluating adaptation and olivocochlear efferent feedback as potential explanations of psychophysical overshoot,” Journal of the Association for Research in Otolaryngology, 12, 345–360. doi:10.1007/s10162-011-0256-5

42. Kawase, T., Delgutte, B., and Liberman, M. C. (1993). “Antimasking effects of the olivocochlear reflex. II. Enhancement of auditory-nerve response to masked tones,” Journal of Neurophysiology, 70, 2533–2549. doi:10.1152/jn.1993.70.6.2533

43. Kishan, A. U., Lee, C. C., and Winer, J. A. (2011). “Patterns of olivocochlear axonal branches,” Open J Neurosci, 1, 2.

44. Kondylidis, K., Vavakou, A., and Van Der Heijden, M. (2025). “The spatial buildup of nonlinear compression in the cochlea,” Front. Cell. Neurosci., 18, 1450115. doi:10.3389/fncel.2024.1450115

45. Liberman, M. C. (1978). “Auditory-nerve response from cats raised in a low-noise chamber,” The Journal of the Acoustical Society of America, 63, 442–455. doi:10.1121/1.381736

46. Liberman, M. C., and Brown, M. C. (1986). “Physiology and anatomy of single olivocochlear neurons in the cat,” Hearing Research, 24, 17–36. doi:10.1016/0378-5955(86)90003-1

47. Lilaonitkul, W., and Guinan, J. J. (2009). “Human Medial Olivocochlear Reflex: Effects as Functions of Contralateral, Ipsilateral, and Bilateral Elicitor Bandwidths,” JARO, 10, 459–470. doi:10.1007/s10162-009-0163-1

48. Lilaonitkul, W., and Guinan, J. J. (2012). “Frequency tuning of medial-olivocochlear-efferent acoustic reflexes in humans as functions of probe frequency,” Journal of Neurophysiology, 107, 1598–1611. doi:10.1152/jn.00549.2011

49. Lopez-Poveda, E. A. (2018). “Olivocochlear Efferents in Animals and Humans: From Anatomy to Clinical Relevance,” Front. Neurol.,, doi: 10.3389/fneur.2018.00197. doi:10.3389/fneur.2018.00197

50. Messing, D. P., Delhorne, L., Bruckert, E., Braida, L. D., and Ghitza, O. (2009). “A non-linear efferent-inspired model of the auditory system; matching human confusions in stationary noise,” Speech Communication, 51, 668–683. doi:10.1016/j.specom.2009.02.002

51. Mogensen, P. K., and Riseth, A. N. (2018). “Optim: A mathematical optimization package for Julia,” JOSS, 3, 1–3. doi:10.21105/joss.00615

52. Osses Vecchi, A., Varnet, L., Carney, L. H., Dau, T., Bruce, I. C., Verhulst, S., and Majdak, P. (2022). “A comparative study of eight human auditory models of monaural processing,” Acta Acust., 6, 17. doi:10.1051/aacus/2022008

53. Rohatgi, A. (n.d.). “WebPlotDigitizer.,”

54. Romero, G. E., and Trussell, L. O. (2021). “Distinct forms of synaptic plasticity during ascending vs descending control of medial olivocochlear efferent neurons,” eLife,, doi: 10.7554/elife.66396. doi:10.7554/elife.66396

55. Romero, G. E., and Trussell, L. O. (2022). “Central circuitry and function of the cochlear efferent systems,” Hearing Research, 425, 108516. doi:10.1016/j.heares.2022.108516

56. Rosowski, J. J. (1991). “The effects of external-and middle-ear filtering on auditory threshold and noise-induced hearing loss,” The Journal of the Acoustical Society of America, 90, 124–135. doi:10.1121/1.401306

57. Roverud, E., and Strickland, E. A. (2015). “Exploring the source of the mid-level hump for intensity discrimination in quiet and the effects of noise,” The Journal of the Acoustical Society of America, 137, 1318–1335. doi:10.1121/1.4908243

58. Ryugo, D. K. (2008). “Projections of low spontaneous rate, high threshold auditory nerve fibers to the small cell cap of the cochlear nucleus in cats,” Neuroscience, 154, 114–126. doi:10.1016/j.neuroscience.2007.10.052

59. Ryugo, D. K., and Fay, R. R. (Eds.) (2010). Auditory and vestibular efferents, Springer New York. doi:10.1007/978-1-4419-7070-1

60. Smalt, C. J., Heinz, M. G., and Strickland, E. A. (2014). “Modeling the Time-Varying and Level-Dependent Effects of the Medial Olivocochlear Reflex in Auditory Nerve Responses,” JARO, 15, 159–173. doi:10.1007/s10162-013-0430-z

61. Steenken, F., Pektaş, A., and Köppl, C. (2024). “Age-related changes in olivocochlear efferent innervation in gerbils,” Front. Synaptic Neurosci., 16, 1422330. doi:10.3389/fnsyn.2024.1422330

62. Strickland, E. A. (2004). “The temporal effect with notched-noise maskers: Analysis in terms of input-output functions,” The Journal of the Acoustical Society of America, 115, 2234– 2245. doi:10.1121/1.1691036

63. Strickland, E. A. (2008). “The relationship between precursor level and the temporal effect,” The Journal of the Acoustical Society of America, 123, 946–954. doi:10.1121/1.2821977

64. Verhulst, S., Altoè, A., and Vasilikov, V. (2018). “Computational modeling of the human auditory periphery: Auditory nerve responses, evoked potentials and hearing loss,” Hearing Research, 360, 55–75. doi:10.1016/j.heares.2017.12.018

65. Walsh, K. P., Pasanen, E. G., and McFadden, D. (2010). “Overshoot measured physiologically and psychophysically in the same human ears,” Hearing Research, 268, 22–37. doi:10.1016/j.heares.2010.04.007

66. Warr, W. B. (1992). “Organization of Olivocochlear Efferent Systems in Mammals,” In D. B. Webster, A. N. Popper, and R. R. Fay (Eds.), The Mammalian Auditory Pathway: Neuroanatomy, Springer Handbook of Auditory Research, Springer Verlag, Vol. 1, pp. 401–448.

67. Warr, W. B., and Guinan, J. J. (1979). “Efferent innervation of the organ of corti: two separate systems,” Brain Research, 173, 152–155. doi:10.1016/0006-8993(79)91104-1

68. Warren, E. H., and Liberman, M. C. (1989a). “Effects of contralateral sound on auditory-nerve responses. I. Contributions of cochlear efferents,” Hearing Research, 37, 89–104. doi:10.1016/0378-5955(89)90032-4

69. Warren, E. H., and Liberman, M. C. (1989b). “Effects of contralateral sound on auditory-nerve responses. II. Dependence on stimulus variables,” Hearing Research, 37, 105–121. doi:10.1016/0378-5955(89)90033-6

70. Wersinger, E., and Fuchs, P. A. (2011). “Modulation of hair cell efferents,” Hearing Research, 279, 1–12. doi:10.1016/j.heares.2010.12.018

71. Wojtczak, M., Beim, J. A., and Oxenham, A. J. (2015). “Exploring the Role of Feedback-Based Auditory Reflexes in Forward Masking by Schroeder-Phase Complexes,” JARO, 16, 81–99. doi:10.1007/s10162-014-0495-3

72. Wojtczak, M., Klang, A. M., and Torunsky, N. T. (2019). “Exploring the role of medial olivocochlear efferents on the detection of amplitude modulation for tones presented in noise,” Journal of the Association for Research in Otolaryngology, 20, 395–413. doi:10.1007/s10162-019-00722-6

73. Yasin, I., Drga, V., and Plack, C. J. (2014). “Effect of Human Auditory Efferent Feedback on Cochlear Gain and Compression,” J. Neurosci., 34, 15319–15326. doi:10.1523/jneurosci.1043-14.2014

74. Yasin, I., Liu, F., Drga, V., Demosthenous, A., and Meddis, R. (2018). “Effect of auditory efferent time-constant duration on speech recognition in noise,” The Journal of the Acoustical Society of America, 143, EL112–EL115. doi:10.1121/1.5023502

75. Ye, Y., Machado, D. G., and Kim, D. O. (2000). “Projection of the marginal shell of the anteroventral cochlear nucleus to olivocochlear neurons in the cat,” J Comp Neurol, 420, 127–138.

76. Zilany, M. S. A., and Bruce, I. C. (2006). “Modeling auditory-nerve responses for high sound pressure levels in the normal and impaired auditory periphery,” The Journal of the Acoustical Society of America, 120, 1446–1466. doi:10.1121/1.2225512

77. Zilany, M. S. A., Bruce, I. C., and Carney, L. H. (2014). “Updated parameters and expanded simulation options for a model of the auditory periphery,” The Journal of the Acoustical Society of America, 135, 283–286. doi:10.1121/1.4837815

78. Zilany, M. S. A., Bruce, I. C., Nelson, P. C., and Carney, L. H. (2009). “A phenomenological model of the synapse between the inner hair cell and auditory nerve: Long-term adaptation with power-law dynamics,” The Journal of the Acoustical Society of America, 126, 2390– 2412. doi:10.1121/1.3238250

