## Supplemental Figure 1 for "A computational model of the mammalian auditory periphery with a multichannel, energy-driven, medial olivocochlear reflex"

SUPPLEMENTARY MATERIALS

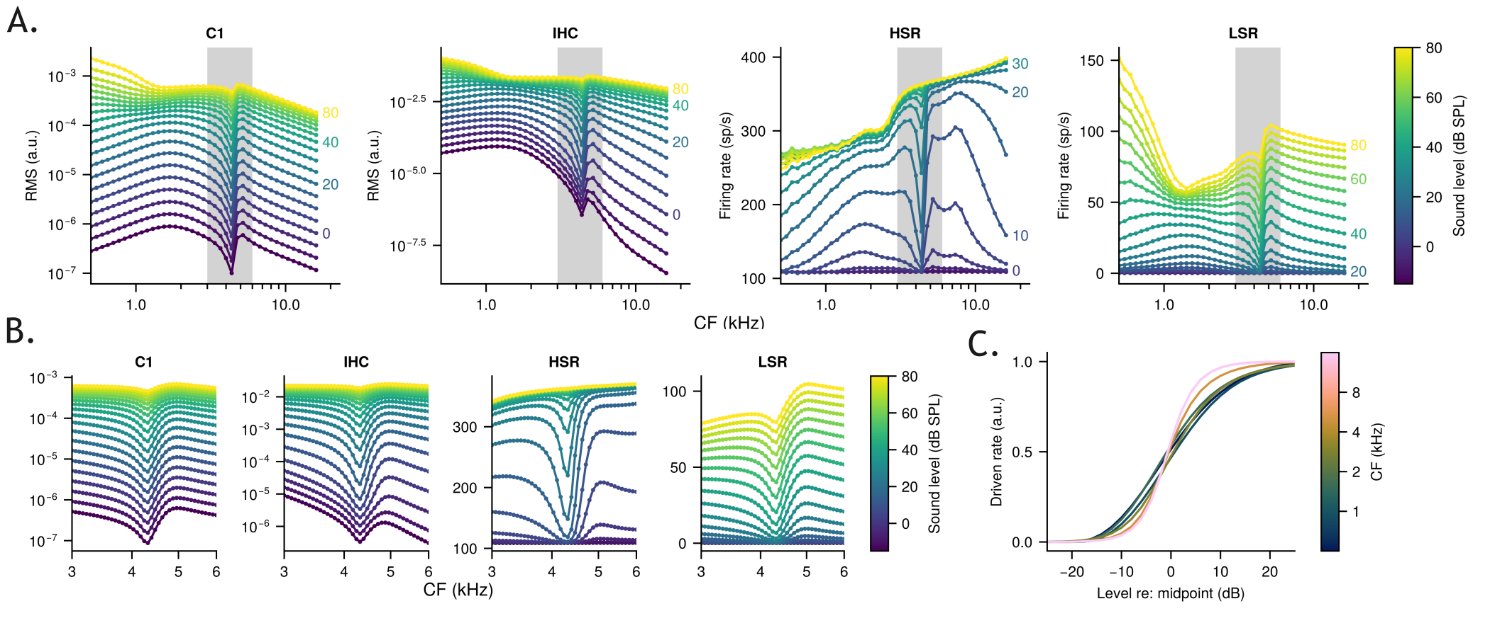

**Supplementary Figure 1.** (A) RMS (C1, IHC) or average (HSR, LSR) responses for an on-CF pure-tone at various sound levels (color) and different CFs (x-axis) in the four stages of the baseline (i.e., without MOC) model. The gray box in the background denotes a region of the CF axis shown in higher resolution in B. (B) Same as A, except over a limited CF range highlighting the effects of the middle ear filter in the model. (C) The HSR results in A except depicted as RLFs normalized to be expressed in terms of driven rate (y-axis) relative to midpoint level (x-axis) rather than absolute level, highlighting that RLFs from higher CFs are steeper (smaller dynamic range) than RLFs at lower CFs.
